# Genome-wide identification of directed gene networks using large-scale population genomics data

**DOI:** 10.1101/221879

**Authors:** René Luijk, Koen F. Dekkers, Maarten van Iterson, Wibowo Arindrarto, Annique Claringbould, Paul Hop, BIOS Consortium, Dorret I. Boomsma, Cornelia M. van Duin, Marleen M.J. van Greevenbroek, Jan H. Veldink, Cisca Wijmenga, Lude Franke, Peter A.C. ’t Hoen, Rick Jansen, Joyce van Meurs, Hailiang Mei, P. Eline Slagboom, Bastiaan T. Heijmans, Erik W. van Zwet

## Abstract

Identification of causal drivers behind regulatory gene networks is crucial in understanding gene function. We developed a method for the large-scale inference of gene-gene interactions in observational population genomics data that are both directed (using local genetic instruments as causal anchors, akin to Mendelian Randomization) and specific (by controlling for linkage disequilibrium and pleiotropy). The analysis of genotype and whole-blood RNA-sequencing data from 3,072 individuals identified 49 genes as drivers of downstream transcriptional changes (*P* < 7 × 10^−10^), among which transcription factors were overrepresented (*P* = 3.3 × 10^−7^). Our analysis suggests new gene functions and targets including for *SENP7* (zinc-finger genes involved in retroviral repression) and *BCL2A1* (novel target genes possibly involved in auditory dysfunction). Our work highlights the utility of population genomics data in deriving directed gene expression networks. A resource of *trans*-effects for all 6,600 genes with a genetic instrument can be explored individually using a web-based browser.

## INTRODUCTION

Identification of the causal drivers underlying regulatory gene networks may yield new insights into gene function^1,2^, possibly leading to the disentanglement of disease mechanisms characterized by transcriptional dysregulation^3^. Gene networks are commonly based on the observed co-expression of genes. However, such networks show only undirected relationships between genes which makes it impossible to pinpoint the causal drivers behind these associations. Adding to this, confounding (e.g. due to demographic and clinical characteristics, technical factors, and batch effects^4–6^) induces spurious correlations between the expression of genes. Correcting for all confounders may prove difficult as some may be unknown^7^. Residual confounding then leads to very large, inter-connected co-expression networks that do not reflect true biological relationships.

To address these issues, we exploited recent developments in data analysis approaches that enable the inference of causal relationships through the assignment of directed gene-gene associations in population-based transcriptome data using genetic instruments^8–10^ (GIs). Analogous to Mendelian Randomization^11,12^ (MR), the use of genetics provides an anchor from where directed associations can be identified. Moreover, GIs are free from any non-genetic confounding. Related efforts have used similar methods to identify novel genes associated with different phenotypes, either using individual level data^8,9^ or using publicly available eQTL and GWAS catalogues^10^. However, these efforts have not systematically taken linkage disequilibrium (LD) and pleiotropy (a genetic locus affecting multiple nearby genes) into account. As both may lead to correlations between GIs, we aimed to improve upon these methods in order to minimize the influence of LD and pleiotropy, and would detect the actual driver genes. This possibly induces non-causal relations^13^, precluding the identification of the specific causal gene involved when not accounted for LD and pleiotropy.

Here, we combine genotype and expression data of 3,072 unrelated individuals from whole blood samples to establish a resource of directed gene networks using genetic variation as an instrument. We use local genetic variation in the population to capture the portion of expression level variation explained by nearby genetic variants (local genetic component) of gene expression levels, successfully identifying a predictive genetic instrument (Gl) for the observed gene expression of 6,600 protein-coding genes. These GIs are then tested for an association with potential target genes *in trans*. Applying a robust genome-wide approach that corrects for linkage disequilibrium and local pleiotropy by modelling nearby GIs as covariates, we identify 49 index genes each influencing up to 33 target genes (Bonferroni correction, *P* < 7 × 10^−10^). Closer inspection of examples reveals that coherent biological processes underlie these associations, and we suggest new gene functions based on these newly identified target genes, e.g. for *SENP7* and *BCL2A1*. An interactive online browser allows researchers to look-up specific genes of interest while using the appropriate, more lenient significance threshold.

## RESULTS

### Establishing directed associations in transcriptome data

We aim to establish a resource of index genes that causally affect the expression of target genes *in trans* using large-scale observational RNA-sequencing data. However, causality cannot be inferred from the correlation between the observed expression measurements of genes, and therefore is traditionally addressed by experimental manipulation. Furthermore, both residual and unknown confounding can induce correlation between genes, possibly yielding to extensive correlation networks that are not driven by biology. To establish causal relations between genes, we assume a structural causal model^14^ describing the relations between genes and using their genetic components, the local genetic variants predicting their expression, as genetic instruments^11^ (GIs). To be able to conclude the presence of a causal effect of the index gene on the target gene, the potential influence of linkage disequilibrium (LD) and pleiotropic effects have to be taken into account, as they may cause GIs of neighbouring genes to be correlated (Figure 1). This is done by blocking the so-called back-door path^14^ from the index GI through the genetic GIs of nearby genes to the target gene by correcting the association between the GI and target gene expression for these other GIs. Note that this path cannot be blocked by adjusting for the observed expression of the nearby genes, as this may introduce collider bias, resulting in spurious associations.

**Figure 1.**
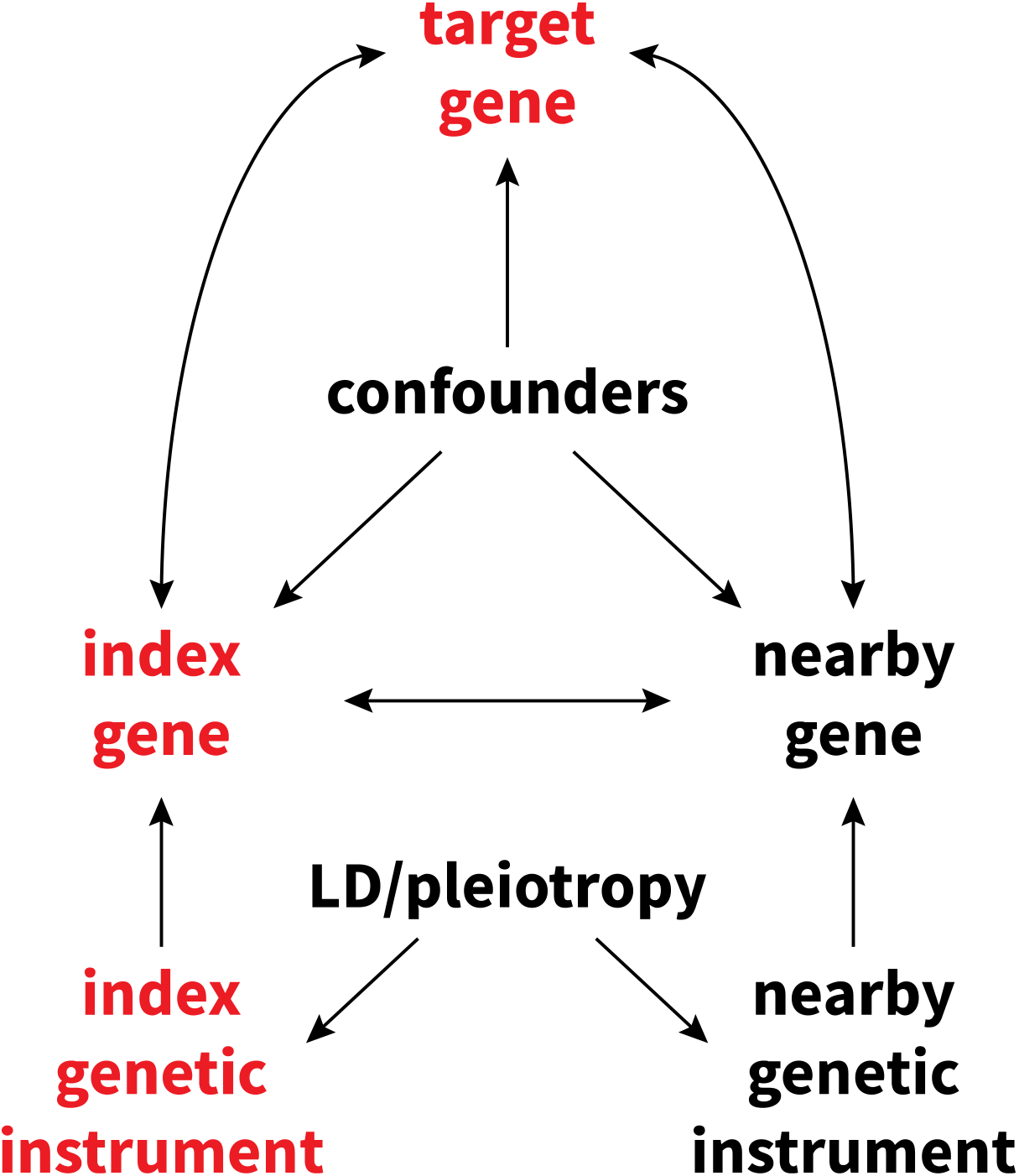
Diagram showing the presumed relations between each variable. A directed arrow indicates the possibility of a causal effect. For instance, the “index genetic instrument” represents nearby SNPs with a possible effect on the nearby gene (analogous to *cis*-eQTLs). A double arrow means the possibility of a causal effect in either direction. The index gene, for example, could have a causal effect on the target gene, or vice versa. We aim to assess the presence of a causal effect of the index gene on the target gene using genetic instruments (GIs) that are free of non-genetic confounding. To do this, we must block the back-door path from the index GI through the GIs of nearby genes to the target gene. This back-door path represents linkage disequilibrium and local pleiotropy and is precluded by correcting for the GIs of nearby genes. Correction for observed gene expression (either of the index gene or of nearby genes) does not block this back-door path, but instead possibly leads to a collider bias, falsely introducing a correlation between the index GI and the target gene.

To assign directed relationships between the expression of genes and establish causality, the first step in our analysis approach was to identify a GI for the expression of each gene, reflecting the local genetic component. To this end, we used data on 3,072 individuals with available genotype and gene expression data (Table S1), measured in whole blood, where we focused on at least moderately expressed (see Methods) protein-coding genes (N = 10,781 genes, Figure S1). Using the 1,021 samples in the training set (see Methods), we obtained a GI consisting of at least 1SNP for the expression of 8,976 genes by applying lasso regression^15^ to nearby genetic variants while controlling for known (cohort, sex, age, cell counts) and unknown covariates^16^ (see Methods). Adding distant genetic variants to the prediction model has been shown to add very little predictive power^8^ and would have induced the risk of including long-range pleiotropic effects.

The strength of the GIs was evaluated using the 2,051 samples in the test set (see Methods). Taking LD and local pleiotropy into account by including the GIs of neighbouring genes (< 1 Mb, Figure 1), we identified 6,600 sufficiently strong GIs having at least partly specific predictive ability (Figure S2A) for the expression its corresponding index gene (*F*-statistic > 10, Figure S1, Table S2). To evaluate the effects of these 6,600 GIs on target gene expression, we used all 3,072 samples to test for an association of each of 6,600 GIs with all of 10,781 expressed, protein-coding genes *in trans* (> 10Mb, Figure S2B). First, this analysis was done without accounting for LD and local pleiotropy (i.e., correcting for neighbouring LD, Figure 1). This genome-wide analysis resulted in 401 directed associations between 134 index genes and 276 target genes after adjustment for multiple testing using the Bonferroni correction (*P* < 7 × 10 ^−10^, Figure 2, Table S3). Among them were 134 index genes affecting the expression of 1 to 33 target genes *in trans* (3.2 genes on average, median of 1 gene), totalling 276 identified target genes. As expected, the resulting networks contained many instances where the same target gene (N = 65) was influenced by multiple neighbouring index genes, hindering the identification of the causal gene. Repeating the analysis for the 134 identified index genes, but corrected for LD and local pleiotropy by including the GIs of neighbouring genes (< 1Mb) resulted in the identification of specific directed effects for 49 index genes on 144 target genes, totalling 156 directed associations (*P* < 7 × 10^−10^, Figure 2), where the number of target genes affected by an index gene varied from 1 to 33 (Table 1, 3.2 genes on average, median of 1 gene). The number of target genes associated with multiple neighbouring index genes drops from 65 to 2, underscoring the importance of correction for LD and local pleiotropy. As this set of 156 directed associations is free from LD and local pleiotropy, and possibly reflect truly causal relations, we use these in further analyses.

**Figure 2.**
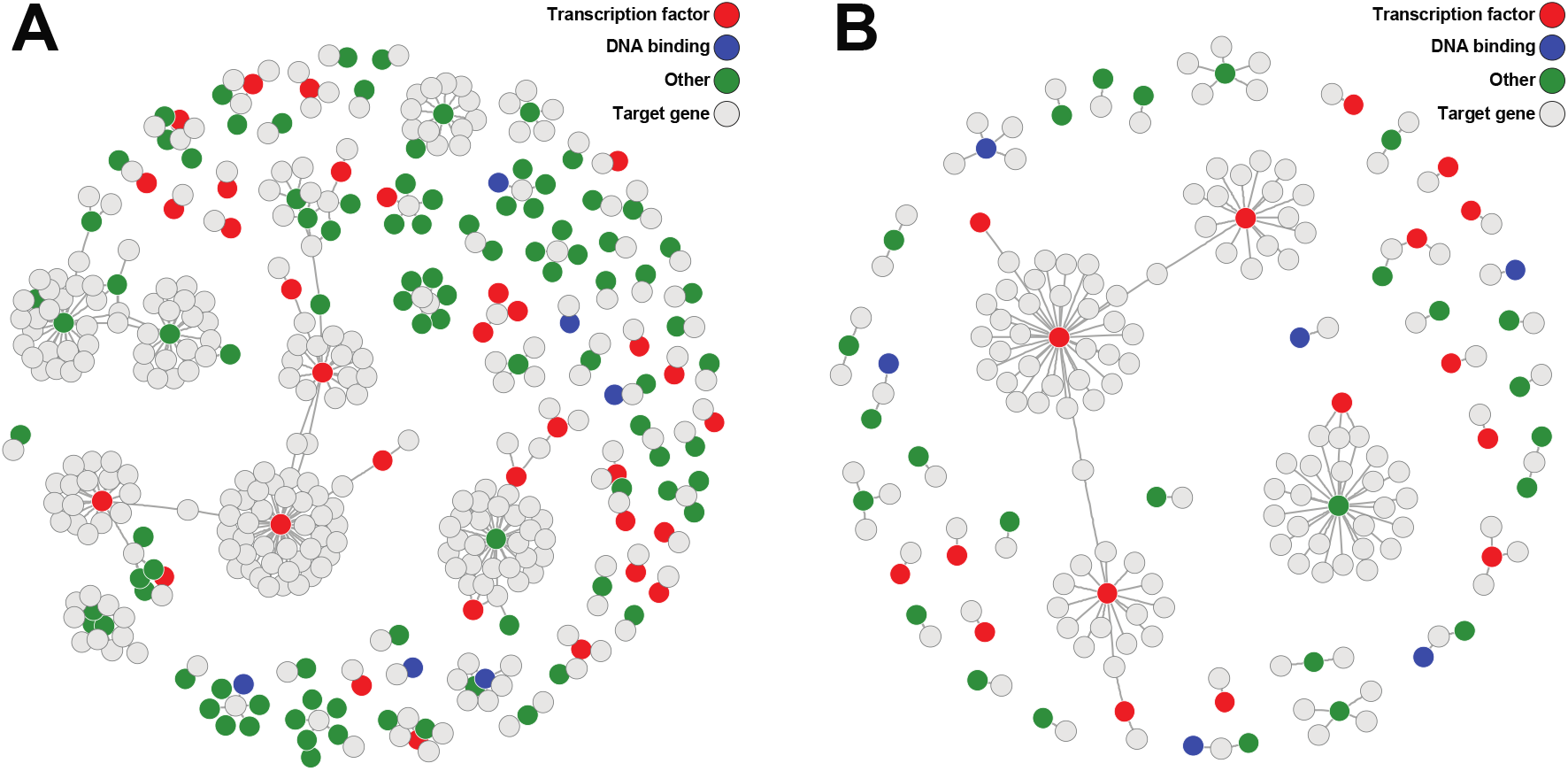
Gene networks showing the directed gene-gene association between genes when not taking LD and local pleiotropy into account (A) and when these are corrected for (B). Index genes identified as a transcription factor are indicated by red circles. Blue circles indicate index genes with DNA binding properties, but are not a known transcription factor^24^. Green circles indicate other index genes. Light grey circles indicate target genes. The uncorrected analysis shows 134 index genes (colored circles) influencing 276 target genes, where several neighbouring index genes seemingly influencing the same target gene, which is reflective of a shared genetic component of those index genes. Specifically, 65 target genes are associated with multiple index genes which lie in close proximity to one another. The number of index genes drop sharply from 134 to 49 (2.7-fold decrease) when do taking LD and local pleiotropy into account. The number of target genes also drops, from 276 to 144 (1.9-fold decrease).

### Validity and stability of the analyses

To ensure the validity and stability of the analyses, we performed several checks regarding common challenges inherent to these analyses and the assumptions underlying them. First, by design, the GIs should be independent of most confounding factors, but confounding may still occur if genetic variants directly affect blood composition, leading to spurious associations. Therefore, we evaluated the association of the 49 GIs with observed red blood cell count and white blood cell counts, and found that none of the 49 GIs were significantly related to any observed cell counts (Figure S3A). In addition, all 156 directed associations remained significant after further adjustment for nearby genetic variants (< 1Mb) reported to influence blood composition^17,18^ (Figure S3B).

To combat any unknown residual confounding and possibly gain statistical power, we added five latent factors to our models, estimated from the observed expression data using cate^16^ (see Methods). We re-tested the 156 identified associations without these factors to evaluate the model sensitivity, showing similar results with slightly attenuated test statistics (Figure S3C). This indicates that our analysis was not influenced by unknown confounding and confirmed the independence of GIs from non-genetic confounding, but did help in reducing the noise in the data, leading to increased statistical power.

Next, to validate the GIs of the 49 index genes, we compared the SNPs constituting the GIs of the 49 index genes associated with target gene expression with previous *cis*-eQTL mapping efforts. While similar sets of genes may be identified using a *cis*-eQTL approach, the utility of using multi-SNP GIs over single-SNP GIs (akin to *cis*-eQTLs) is shown in the increased predictive ability of multi-SNP GIs (Figure S3D), and thus in the number of predictive GIs. Only 4,910 single-SNP GIs were predictive of its corresponding index gene (*F*-statistic > 10), compared to 6,600 multi-SNP instrumental variables. Of the 49 index genes corresponding to the 49 GIs, 47 genes (96.1%) were previously identified as harbouring a c/s-eQTL in large subset of the whole blood transcriptome data we analysed here (N = 2,116), using an independent analysis strategy^19^. Almost all of the corresponding GIs (98%, N = 46) were strongly correlated with the corresponding eQTL SNPs (R2 > 0.8). Similarly, 26 of the 49 index genes (53%) were also reported as having a *cis*-eQTL effect in a much smaller set of whole blood samples (N = 338) part of GTEx^20^, 23 of which also correlated strongly with the corresponding eQTL-SNPs (R2 > 0.8). When considering all tissues in the GTEx project, we found 48 of 49 index genes were identified as harbouring a *cis*-eQTL in any of the 44 tissues measured.

Next, we compared our identified effects with *trans*-eQJls identified earlier in whole-blood samples^21^. First, we found 97 target genes identified here (67%) overlapped with those found by Joehanes *et al*., 19 of which had their corresponding GI and lead SNP in close proximity (< 1Mb, Figure S4), suggesting that the effects are indeed mediated by the index gene assigned using our approach. Testing for a *cis*-eQTL of those SNPs identified by Joehanes *et al*. on the nearby index genes, we found all 19 index genes indeed had at least one nearby lead SNP that influenced its expression (*P* < 6 × 10^−4^, Table S4). This number increased to 31 at a look-up threshold for multiple testing in our analysis (*P* < 4.6 × 10^−6^), indicating that limited statistical power of both studies may lead to an underestimation of the overlap.

As a last check, we investigated potential mediation effects of each of the 49 GIs by observed index gene expression (Figure 1), meaning the effect of a GI on target gene expression should diminish when correcting for the observed index gene expression. However, small effect sizes and considerable noise in both mediator and outcome lead to low statistical power to detect mediated effects^22,23^. Nevertheless, we found 105 of 156 significant directed associations (67%) to show evidence for mediation (Bonferroni correction: *P* < 0.00031; Table S5).

### Exploration of directed networks

To gain insight in the molecular function of 49 index genes affecting target gene expression, we used Gene Ontology (GO) to annotate our findings. The set of 49 index genes was overrepresented in the GO terms DNA Binding (*P* = 5 × 10^−8^) and Nucleic Acid Binding (*P* = 2.8 × 10^−5^, Table S6), with 43.8% (N = 21) and 47.9% (N = 23) of genes overlapping with those gene sets, respectively. In line with this finding, we found a significant overrepresentation of transcription factors (N = 17; odds ratio = 5.7, *P* = 3.3 × 10^−7^) using a manually curated database of transcription factors^24^. We note that such enrichments are expected a priori and hence indirectly validate our approach. Of interest, several target genes of two transcription factors overlapped with those identified in previous studies^25,26^ (*IKZF1*: 27% of its target genes, N = 4; *PLAGL1*:15% of its target genes, N = 5). Using a more lenient significance threshold corresponding to a look-up for each of these 17 transcription factors (thus correcting for only 10,781 potential target genes; *P* < 4.6 × 10^−6^), we identified overlapping target genes for an additional 3 transcription factors^25–28^ (*CREB5, NFKB1, NKX3-1*) and a total of 38 TF-target gene pairs corresponding between our analysis and previous studies (Table S7).

Finally, we explore the biological processes that are revealed by our analysis for several index genes that either are known transcription factors^24^ or affect many genes in *trans*. While these results are limited to Bonferroni-significant directed associations (*P* < 7 × 10^−10^, correcting for all possible combinations of the 6,600 index genes and 10,781 target genes), researchers can interactively explore the whole resource by means of a look-up at a much more lenient significance threshold (*P* < 2.9 × 10^−6^, testing for a gene to have an effect *in trans*, or being affected by other genes, totalling 17,381 tests (6,600 + 10,781)) using a dedicated browser (see URLs).

#### Sentrin/SUMO-specific proteases 7 (SENP7)

We identified 25 target genes to be affected *in trans* by sentrin/small ubiquitin-like modifier (SUMO)-specific proteases 7 (*SENP7*, Figure 3, Figure 4, Table 1), significantly expanding on the five previously suspected target genes resulting from an earlier expression QTL approach^29^. Increased *SENP7* expression resulted in the upregulation of all but one of the target genes (96%). Remarkably, 23 of the 25 target genes affected by *SENP7* are zinc finger protein (ZFP) genes located on chromosome 19. More specifically, 18 target genes are located in a 1.5Mb ZFP cluster mapping to 19q13.43 (Figure 3). ZFPs in this cluster are known transcriptional repressors, particularly involved in the repression of endogenous retroviruses^30^. Parallel to this, *SENP7* has also been identified to promote chromatin relaxation for homologous recombination DNA repair, specifically through interaction with chromatin repressive KRAB-Association Protein (*KAP1*, also known as *TRIM28*). *KAP1* had already been implicated in transcriptional repression, especially in epigenetic repression and retroviral silencing^31,32^, although *KAP1* had no predictive GI (*F*-statistic = 4.9). Therefore, it has been speculated *SENP7* may also play a role in retroviral silencing^33^. Given the widespread effects of *SENP7* on the transcription of chromosome 19-linked ZFPs involved in retroviral repression^30^, it corroborates a role of *SENP7* in the repression of retroviruses, specifically through regulation of this ZFP cluster. *SENP7* is not a TF and does not bind DNA, but considering it is a SUMOylation enzyme, it possibly has its effect on the ZFP cluster through deSUMOylation of *KAP1*^34^.

**Figure 3.**
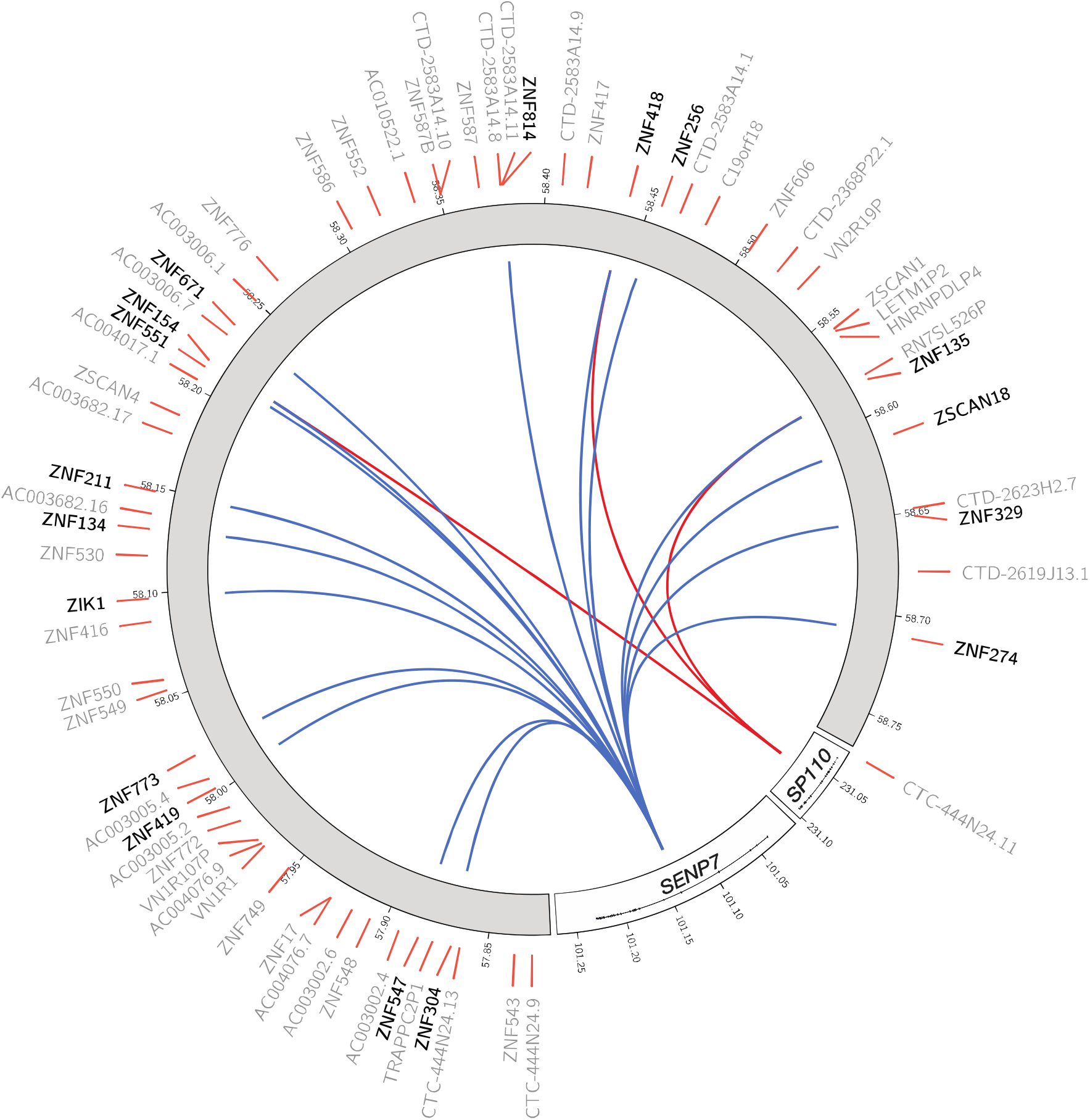
*SENP7* (chromosome 3) and *SP110* (chromosome 2) affect a zinc finger cluster located on chromosome 19 involved in retroviral repression, among others. Blue lines indicate a positive association (upregulation), red lines indicate a negative association (downregulation). Colouring indicates consistent opposite effects of *SENP7* and *SP110* on their shared target genes.

#### SP110 nuclear body protein (SP110)

In our genome-wide analysis, we found that the transcription factor *SP110* nuclear body protein *(SP110)* influences three zinc finger proteins (Figure 3, Figure 4). During viral infections in humans, *SP110* has been shown to interact with the Remodelling and Spacing Factor 1 (*RSF1*) and Activating Transcription Factor 7 Interacting Protein (*ATF7IP*), suggesting it is involved in chromatin remodelling^35^. Interestingly, all three of the genes targeted by *SP110* are also independently influenced by *SENP7*, although *SP110* shows opposite effects (Figure S5), and are located in the same ZFP gene cluster on chromosome 19. A specific look-up (thus relaxing the multiple testing burden; Figure 3b) for *SP110* targets show six genes, all also independently affected by *SENP7*. This overlap of target genes supports the previous suggestion that *SP110* is involved in the innate antiviral response^36^, presumably through regulation of the same ZPF cluster regulated by *SENP7*.

**Figure 4.**
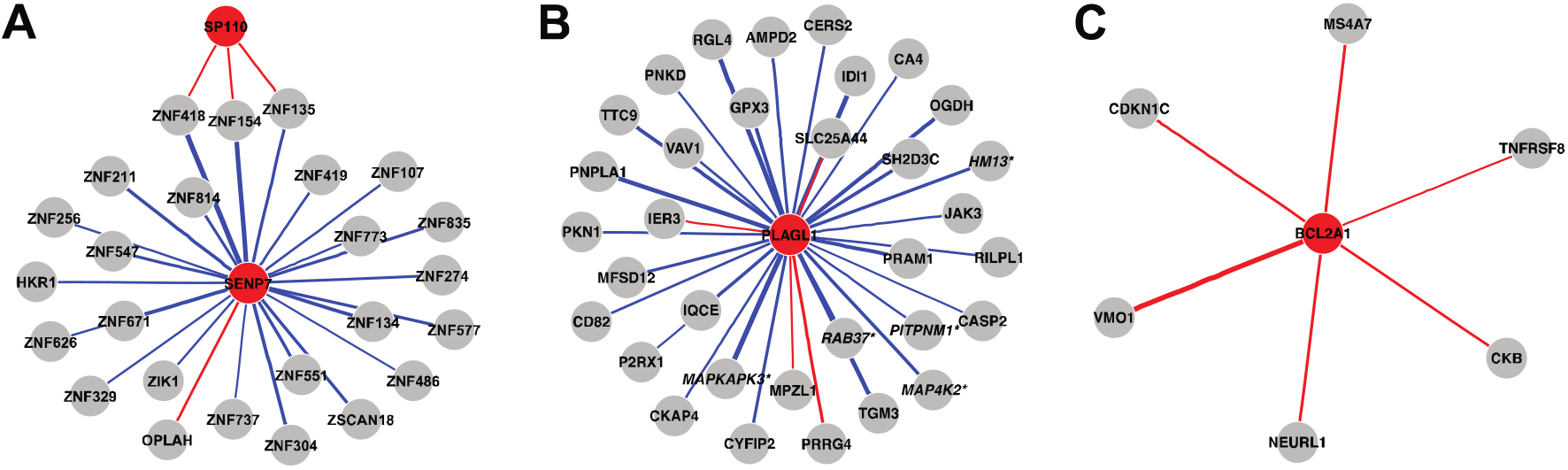
Identified target genes for *SENP7* (A), *SP110* (A), *PLAGL1* (B), and *BCL2A1* (C). Starred and italic gene names indicate previously reported target genes^25–28^. Blue and red lines indicate positive and negative associations, respectively; line thickness indicates strength of the association.

#### Pleiomorphic adenoma gene-like 1 (PLAGL1)

The index gene with the most identified target gene effects *in trans* is Pleiomorphic Adenoma Gene-Like 1 (*PLAGL1*, also known as *LOT1, ZAC). PLAGL1* is a transcription factor and affected 33 genes, 29 of which are positively associated with *PLAGL1* expression (88%, Figure 4). *PLAGL1* is part of the imprinted *HYMAI/ZAC1* locus, which has a crucial role in fetal development and metabolism^37,38^. This locus, and overexpression of *PLAGL1* specifically, has been associated with transient neonatal diabetes mellitus^35,39^ (TNDM) possibly by reducing insulin secretion^40^. *PLAGL1* is known to be a transcriptional regulator of PACAP-type I receptor^41^ (*PAC1-R). PACAP*, in turn, is a regulator of insulin secretion^42,43^. In line with these findings, we found several target genes to be involved in metabolic processes. Most notably, we identified *MAPKAPK3* (MK3) and *MAP4K2* to be upregulated by *PLAGL1*, previously identified as *PLAGL1* targets^28^, and both part of the mitogen-activated protein kinase (MAPK) pathway. This pathway has been observed to be upregulated in type II diabetic patients (reviewed in ^44^). In addition, inhibition of *MAPKAP2* and *MAPKAP3* in obese, insulin-resistant mice has been shown to result in improved metabolism^45^, in line with the association between upregulation of *PLAGL1* and the development of TNDM. Furthermore, *PLAGL1* may be implicated in lipid metabolism and obesity through its effect on *IDI1, PNPLA1, JAK3*, and *RAB37* expression^46–49^. While not previously established as target genes, they are in line with the proposed role of *PLAGL1* in metabolism^37,38^.

#### Bcl-related protein A1 (BCL2A1)

Increased expression of Bcl-related protein A1 (*BCL2A1*) downregulated all five identified target genes (Figure 4). *BCL2A1* encodes a protein part of the B-cell lymphoma 2 (*BCL2*) family, an important family of apoptosis regulators. It has been implicated in the development of cancer, possibly through the inhibition of apoptosis (reviewed in^50^). One target gene, *NEURL1*, is known to cause apoptosis^51^, in line with its strong negative association with *BCL2A1* expression. Similarly, *CDKN1C* was also downregulated by *BCL2A1*, and implicated in the promotion of cell death^52–55^. However, little is known about the strongest associated target gene, *VMO1* (*P* = 1.5 × 10^−8^). It has been implicated in hearing, due to its highly abundant expression in the mouse inner ear^56^, where *BCL2A1* may have a role in the development of hearing loss through apoptosis, since cell death is a known contributor to hearing loss in mice^57^. In line with its role in the inhibition of apoptosis, *BCL2A1* overexpression has a protective effect on inner ear mechanosensory hair cell death in mice^58^. Lastly, the target gene *CKB* has also been implicated in hearing impairment in mice^59^ and Huntington’s disease^60^, further suggesting a role of *BCL2A1* in auditory dysfunction.

#### Mediation of target gene expression through local DNA methylation

Previously, genetic variants have been found to influence DNA methylation *in trans*^29,61^. Methylation, in turn, can have a causal effect on gene expression (discussed in^62^). This led us to hypothesize that the directed effects on target gene expression identified here could be mediated by changes in DNA methylation near those target genes. We investigated this hypothesis by first obtaining a single score per target gene by summarizing the methylation of nearby CpGs, similar to the construction of the GIs (see Methods), reflective of the local methylation landscape of the target gene. Next, we globally tested for mediation of the identified effects by the methylation scores using Sobel’s test^63^. Evidence for mediation by local changes in DNA methylation were found for 33 effects, pertaining to 8 index genes and 31 target genes (Table S8). Most notably, the mediation analysis showed most of the *SENP7* effects on target gene expression are mediated by local changes in methylation (22 genes, 88%). To further investigate which CpGs specifically are responsible for mediating those 33 effects, we tested each CpG constituting the methylation scores separately, identifying 95 CpGs. Most of the 95 CpGs lie adjacent to a CpG island (CGI), in so-called CGI shores^64,65^ (N = 41, OR = 2.9, *P* = 1.3 × 10^−5^). This suggests regulation of several target genes is at least partly mediated by local changes in DNA methylation or correlated epigenomic markers.

## DISCUSSION

In this work, we report on an approach that uses population genomics data to generate a resource of directed gene networks. Our genome-wide analysis of whole-blood transcriptomes yields strong evidence for 49 index genes to specifically affect the expression of up to 33 target genes *in trans*. We suggest previously unknown functions of several index genes based on the identification of new target genes. Researchers can fully exploit the utility of the resource to look up *trans*-effects of a gene of interest using an interactive gene network browser while using an appropriate, more lenient significance threshold, instead of the strict significance threshold used in our genome-wide analysis.

The identified directed associations provide novel mechanistic insight into gene function. Many of the 49 index genes affecting target gene expression are established transcription factors (TFs), or are known for having DNA binding properties, an anticipated observation supporting the validity of our analysis. The identification of non-TFs will in part relate to the fact that the effect of an index gene may regulate the activity of TFs, for example by post-translational modification. This is illustrated by *SENP7* that we observed to concertedly affect the expression of zinc finger protein genes involved in the repression of retroviruses, likely by deSUMOylation of the transcription factor *KAP1*^34^. Other mechanistic insights that can be distilled from these results include the potential involvement of *BCL2A1* in auditory dysfunction, conceivably through the regulation of apoptosis.

While observational gene expression data can be used to construct gene co-expression networks^60^, which is sometimes complemented with additional experimental information^28^, such an approach lacks the ability to assign causal directions. Experimental approaches using CRISPR-cas9 coupled with single-cell technology^66–68^ are in principle able to demonstrate causality at a large scale, but only in vitro, while the advantage of observational data is that it reflects in vivo situations. These experimental approaches currently rely on extensive processing of single-cell data that is associated with high technical variability^66^, complicating the construction of specific gene-gene associations. In addition, off-target effects of CRISPR-cas9 cannot be excluded^69^, potentially influencing the interpretation of these experiments. Finally, such efforts are currently limited in the number of genes tested^66–68^, whereas we were able to perform a genome-wide analysis. Hence, experimental and population genomics approaches are complementary in identifying causal gene networks.

Traditional *trans*-eQTL studies aim to find specific genetic loci associated with distal changes in gene expression^21,70^. The limitation of this approach is that they are not designed to assign the specific causal gene responsible for the trans-effect because they do not control for LD and local pleiotropy (a genetic locus affecting multiple nearby genes). Hence, our approach enriches trans-eQTL approaches by specifying which index gene induces changes in target gene expression. However, it does not detect *trans*-effects independent of effects on local gene expression. In addition, identification of the causal path using a trans-eQTL approach is difficult to establish. Testing for mediation through local changes in expression^23,71^ may be limited in statistical power, as these approaches are designed to only test the mediation effect of one lead SNP^23^.

The application of related analysis methods was recently used to infer associations between gene expression and phenotypic outcomes (instead of gene expression as we did here). Two studies first constructed multi-marker GIs in relatively small sample sets to then apply these GIs in large datasets without gene expression data^8,9^. A different, summary-data-based Mendelian randomization (SMR) approach identifies genes associated with complex traits based on publicly available GWAS and eQTL catalogues^10^. However, neither of these approaches take LD and pleiotropic effects into account, led to many neighbouring, nonspecific effects^8–10^. We show that correcting for these LD and local pleiotropy will aid in the identification of the causal gene, as opposed to the identification of multiple, neighbouring genes, analogous to fine mapping in GWAS. Furthermore, the use of eQTL and GWAS catalogues are usually the result of genome-wide analyses, where only statistically significant variants are taken into account. Here, we use the full genetic landscape surrounding a gene, thereby maximizing the predictive ability of expression measurements by our GIs^8^. While we have used our genome-wide approach to identify directed gene networks, we note this method may also be used to annotate trait-associated variants with potential target genes, either by using individual level data^8,9^, or by using SMR^10^.

The analysis approach presented here relies on using GIs of expression of an index gene as a causal anchor, an approach akin to Mendelian randomization^11^. While GIs could provide directionality to bi-directional associations in observational data, genetic variation generally explains a relatively small proportion of the variation in expression (Figure S2A). The GIs for index gene expression identified here are no exception, significantly limiting statistical power of similar approaches^72,73^. Increased sample sizes and improvement on the prediction of index gene expression will help in identifying more target genes.

Our current analysis strategy aims for causal inference, obviating LD and local pleiotropic effect by correcting for the GIs of nearby genes. However, we only corrected for GIs of genes within 1Mb of the current index gene, leaving the possibility of pleiotropic effects beyond this threshold. For example, the GI of an index gene may influence both the expression of the index gene and another gene, located outside of the 1Mb window, where the induced changes in that genes’ expression are the causal factor of the identified target genes. A related problem arises when a shared genetic component between neighbouring index genes causes all of them to associate with a single distant target gene, hindering the identification of the index gene responsible for the induced trans-effect. By correcting for the GI of nearby genes, these potentially biologically relevant effects are lost (Figure 1).

As many genetic variants have been shown to affect methylation *in trans*^29,61^, we hypothesized that the identified *trans*-effects here may be mediated by target gene methylation. A limited number of directed associations show evidence for mediation by target gene methylation. This is in line with earlier observations regarding a limited overlap between eQTLs and meQTLs^61^, and suggests changes in transcriptional activity may not always be reflected by altered methylation levels^74^. Alternatively, long-range effects^75^, or other, uncorrelated epigenetic processes could act as a mediator. Furthermore, a bidirectional interplay between DNA methylation and gene expression possibly makes their relationship more intricate than previously appreciated^71^.

In conclusion, we present a genome-wide approach that identifies causal effects of gene expression on distal transcriptional activity in population genomics data and showcase several examples providing new biological insights. The resulting resource is available as an interactive network browser that can be utilized by researchers for look-ups of specific genes of interest (see URLs).

## Methods

### Cohorts

The Biobank-based Integrative Omics Study (BIOS, Additional SI1) Consortium comprises six Dutch biobanks: Cohort on Diabetes and Atherosclerosis Maastricht^76^ (CODAM), LifeLines-DEEP^77^ (LLD), Leiden Longevity Study^78^ (LLS), Netherlands Twin Registry^79,80^ (NTR), Rotterdam Study^81^ (RS), Prospective ALS Study Netherlands^82^ (PAN). The data that were analysed in this study came from 3,072 unrelated individuals (Supplementary Table 1). Genotype data, DNA methylation data, and gene expression data were measured in whole blood for all samples. In addition, sex, age, and cell counts were obtained from the contributing cohorts. The Human Genotyping facility (HugeF, Erasmus MC, Rotterdam, The Netherlands, http://www.blimdna.org) generated the methylation and RNA-sequencing data.

### Genotype data

Genotype data were generated within each cohort. Details on the genotyping and quality control methods have previously been detailed elsewhere (LLD: Tigchelaaret et al.^77^; LLS: Deelen et a1.^83^; NTR: Lin et al.^84^; RS: Hofman et al.^81^; PAN: Huisman et al.^82^.

For each cohort, the genotype data were harmonized towards the Genome of the Netherlands^85^ (GoNL) using Genotype Harmonizer^86^ and subsequently imputed per cohort using Impute2^87^ and the GoNL reference panel^85^ (v5). We removed SNPs with an imputation info-score below 0.5, a HWE *P* < 10^−4^, a call rate below 95% or a minor allele frequency smaller than 0.01. These imputation and filtering steps resulted in 7,545,443 SNPs that passed quality control in each of the datasets.

### Gene expression data

A detailed description regarding generation and processing of the gene expression data can be found elsewhere^19^. Briefly, total RNA from whole blood was deprived of globin using Ambion’s GLOBIN clear kit and subsequently processed for sequencing using Illumina’s Truseq version 2 library preparation kit. Paired-end sequencing of 2x50bp was performed using lllumina’s Hiseq2000, pooling 10 samples per lane. Finally, read sets per sample were generated using CASAVA, retaining only reads passing Illumina’s Chastity Filter for further processing. Data were generated by the Human Genotyping facility (HugeF) of ErasmusMC (The Netherlands, see URLs). Initial QC was performed using FastQC (v0.10.1), removal of adaptors was performed using cutadapt^88^ (v1.1), and Sickle^89^ (v1.2) was used to trim low quality ends of the reads (minimum length 25, minimum quality 20). The sequencing reads were mapped to human genome (HG19) using STAR^90^ (v2.3.0e).

To avoid reference mapping bias, all GoNL SNPs (http://www.nlgenome.nl/?page_id=9) with MAF > 0.01 in the reference genome were masked with N. Read pairs with at most 8 mismatches, mapping to as most 5 positions, were used.

Gene expression quantification was determined using base counts^19^. The gene definitions used for quantification were based on Ensembl version 71, with the extension that regions with overlapping exons were treated as separate genes and reads mapping within these overlapping parts did not count towards expression of the normal genes.

For data analysis, we used counts per million (CPM), and only used protein coding genes with sufficient expression levels (median log(CPM) > 0), resulting in a set of 10,781 genes. To limit the influence of any outliers still present in the data, the data were transformed using a rank-based inverse normal transformation within each cohort.

### DNA methylation data

The Zymo EZ DNA methylation kit (Zymo Research, Irvine, CA, USA) was used to bisulfite-convert 500 ng of genomic DNA, and 4 μl of bisulfite-converted DNA was measured on the Illumina HumanMethylation450 array using the manufacturer’s protocol (Illumina, San Diego, CA, USA). Preprocessing and normalization of the data were done as described earlier^91^. In brief, IDAT files were read using the *minfi* R package^92^, while quality control (QC) was performed using *MethylAid*^93^. Filtering of individual measurements was based on detection P-value (*P* < 0.01), number of beads available (≤ 2) or zero values for signal intensity, followed by the removal of ambiguously mapped probes^94^. Normalization was done using Functional Normalization^95^ as implemented in the *minfi* R package^92^, using five principal components extracted using the control probes for normalization. All samples or probes with more than 5% of their values missing were removed. The final dataset consisted of 440,825 probes measured in 3,072 samples. Similar to the RNA-sequencing data, we also transformed methylation data using a rank-based inverse normal transformation within each cohort, to limit the influence of any remaining outliers.

### Constructing a local genetic instrumental variable for gene expression

We started by constructing genetic instruments (GIs) for the expression of each gene in our data. We first split up the genotype and RNA-sequencing data in a training set (one-third of all samples, N = 1,021) and a test set (two-thirds of all samples, N = 2,051), making sure all cohorts and both sexes were evenly distributed over the train and test sets (57% female), as well as an even distribution of age (mean = 56, sd = 14.8). Using the training set only, we built a GI for each gene separately that best predicts its expression levels using lasso^15^, using nearby genetic variants only (either within the gene or within 100kb of a gene’s TSS or TES), while correcting for both known (cohort, sex, age, cell counts) and unknown covariates. Estimation of the unknown covariates was done by applying *cate*^16^ to the observed expression data, leading to 5 unknown latent factors used. Those factors, together with the known covariates, were left unpenalised. To estimate the optimal penalization parameter *λ*, we used five-fold cross-validation as implemented in the R package *glmnet*^96^. The obtained GI consists of a weighted linear combination of the individual dosage values, weighted by the shrunken regression coefficients, yielding one value per individual for each GI. We then evaluated its predictive ability in the test set by employing Analysis of Variance (ANOVA) to evaluate the added predictive power of the GI over the covariates and neighbouring GIs (within 1Mb), as reflected by the *F*-statistic (*F* > 10).

### Testing for *trans*-effects

Using linear regression, we tested for an association between each GI and the expression of potential target genes *in trans* (> 10Mb), while correcting for known (cohort, sex, age, cell counts) and unknown covariates, as well as GIs of nearby genes (< 1Mb). Missing observations in the measured red blood cell count (RBC) and white blood cell counts (WBC) were imputed using the R package *pls*, as described earlier^6^. Any inflation or bias in the test-statistics was estimated and corrected for using the R package *bacon*^6^. Correction for multiple testing was done using Bonferroni (*P* < 10^−10^). The resulting networks were visualized using the R packages *network* and *ndtv*.

### Mediation analysis

To identify CpGs mediating the effect of the genetic instrumental variable (GI) on the target gene, we first summarised the local methylation landscape around each target gene using a method similar to the creation of the GIs. We used lasso to predict target gene expression based on all nearby CpGs in the train set (either located in the target gene or within 250 Kb), using five-fold cross-validation to optimize the penalization parameter *λ*. This resulted in one score reflecting this methylation landscape, whose predictive ability of the target gene’s expression we assessed using ANOVA in the test set (*F* > 10).

In order to assess the mediation of the GI on its target gene through DNA methylation, we employed the Sobel test^63^. This method is based on the notion that the influence of an independent variable (the GI) on a dependent variable (expression of the target gene) should diminish, or even disappear, when controlling for a mediator (methylation score).

### Enrichment analyses

Functional analysis of gene sets was performed for GO Molecular Function annotations using DAVID^97^, providing a custom background consisting of all genes with a predictive GI (*F* > 10). Fisher’s exact test was employed to specifically test for an enrichment of transcription factors using manually curated database of transcription factors^24^.

### URLs

Look-ups can be performed using our interactive gene network browser at http://bios-vm.bbmrirp3-lumc.surf-hosted.nl:8008/NetworkBrowser/. Data were generated by the Human Genotyping facility (HugeF) of ErasmusMC, the Netherlands (http://www.glimDNA.org). Webpages of participating cohorts: LifeLines, http://lifelines.nl/lifelines-research/general; Leiden Longevity Study, http://www.healthy-ageing.nl/ and http://www.leidenlangleven.nl/; Netherlands Twin Registry, http://www.tweelingenregister.org/; Rotterdam Studies, http://www.erasmusmc.nl/epi/research/The-Rotterdam-Study/; Genetic Research in Isolated Populations program, http://www.epib.nI/research/geneticepi/research.html#gip; CODAM study, http://www.carimmaastricht.nl/; PAN study, http://www.alsonderzoek.nl/.

### Accession codes

Raw data were submitted to the European Genome-phenome Archive (EGA) under accession EGAS00001001077.

## Acknowledgments

This research was financially supported by BBMRI-NL, a Research Infrastructure financed by the Dutch government (NWO, numbers 184.021.007 and 184.033.111). Samples were contributed by LifeLines, the Leiden Longevity Study, the Netherlands Twin Registry (NTR), the Rotterdam Study, the Genetic Research in Isolated Populations program, the Cohort on Diabetes and Atherosclerosis Maastricht (CODAM) study and the Prospective ALS study Netherlands (PAN). We thank the participants of all aforementioned biobanks and acknowledge the contributions of the investigators to this study. This work was carried out on the Dutch national e-infrastructure with the support of SURF Cooperative. We acknowledge the support from the Netherlands Cardiovascular Research Initiative (the Dutch Heart Foundation, Dutch Federation of University Medical Centres, the Netherlands Organisation for Health Research and Development, and the Royal Netherlands Academy of Sciences) for the GENIUS project “Generating the best evidence-based pharmaceutical targets for atherosclerosis” (CVON2011-19).

## Author contributions

Conceptualization, BTH, EWvZ, RL, KFD, MvI; Methodology, RL, WEvZ, MvI; Formal Analysis, RL; Resources, WA, AC, DIB, CMvD, MMJvG, JHV, CW, LF, PACtH, RJ, JvM, HM, PES; Writing - Original Draft, RL; Writing – Review & Editing, RL, BTH, EWvZ, PH AC, DIB, CMvD, MMJvG, JHV, CW, PACtH, RJ, JvM, HM, PES; Visualization, RL, BTH; Supervision, BTH, EWvZ

---

**Figure S1.**
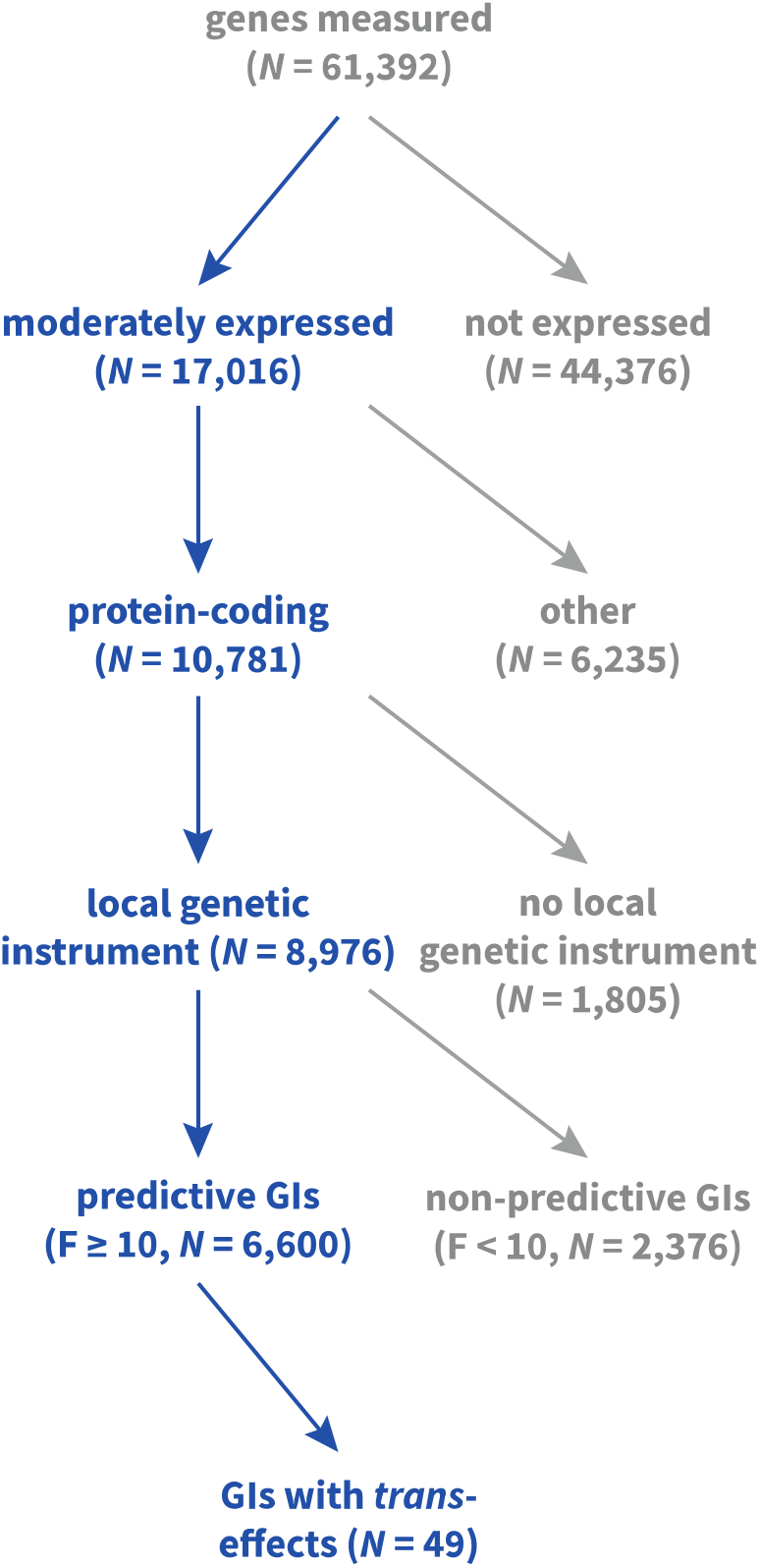
Diagram showing the number of genes and genetic instruments (GIs) in each stage of the analysis.

**Figure S2.**
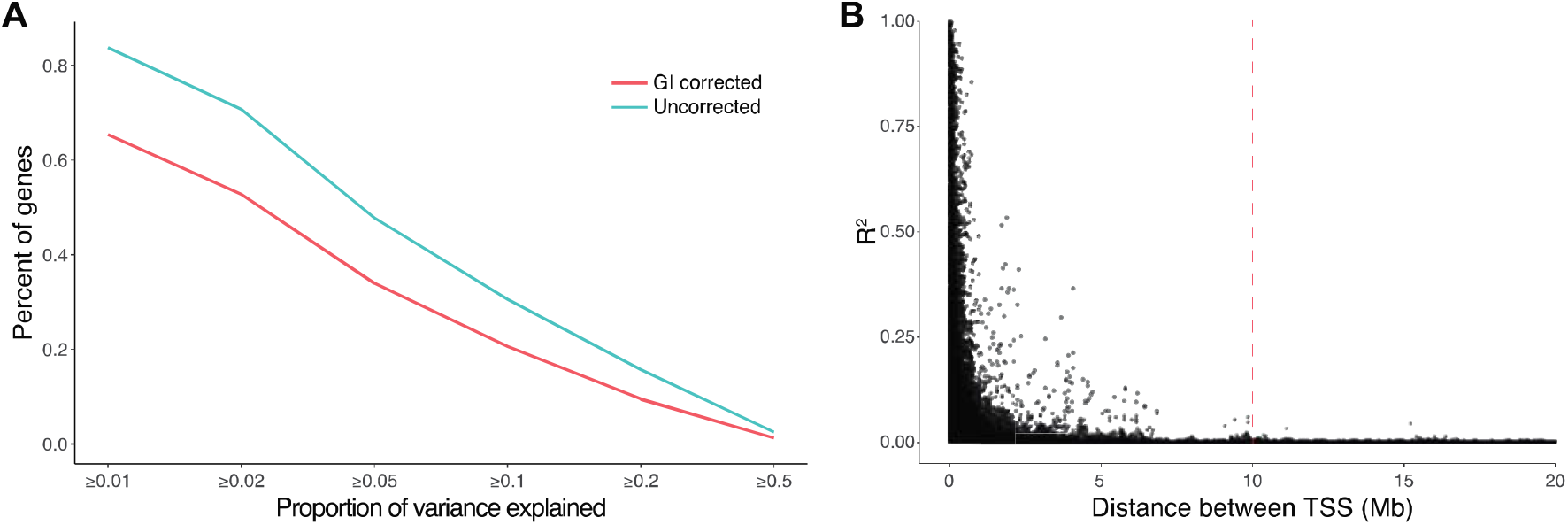
Genetic instruments (GIs) account for a moderate amount of index gene expression variation explained, and are strongly correlated over small distances. A) The proportion of variance (R^2^, x-axis) in index gene expression explained by the corresponding genetic instrumental variable (GI). The blue line indicates the uncorrected R^2^, or the total variance explained by the GI. The red line indicates the R^2^corrected for the GIs of neighbouring index genes, or the proportion of variance explained specifically by the current GI. The proportion of variance explained generally is fairly modest. B) The correlation between genetic instruments (GIs, y-axis) of different genes strongly decreases as the distance (x-axis) between the corresponding genes increases. The median R^2^ between any two GIs corresponding to genes located at least 10Mb (definition of trans, indicated by red dotted line) away from each other is 1.5 × 10^−4^.

**Figure S3.**
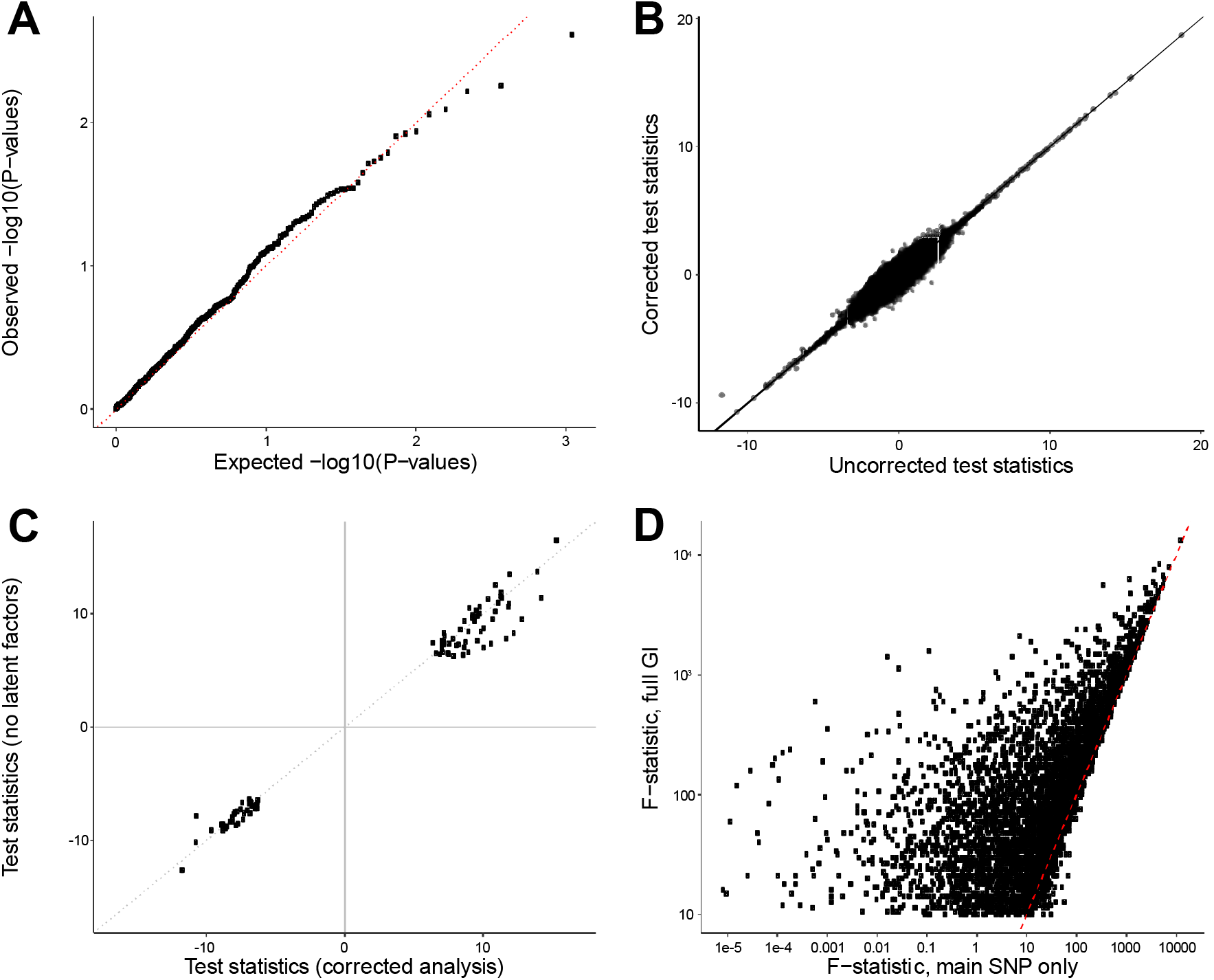
Several checks indicate the stability of our analysis. A) Quantile-quantile plot of the expected −log_10_(*P*-values) (x-axis) and observed −log_10_(*P*-values) (y-axis) resulting from associating all GIs with known cell counts. The observed P-values follow the distribution expected under the null hypothesis, indicative of no association between the GIs and known cell counts. B) All 156 directed associations remained after further adjustment for nearby genetic variants (< 1Mb) reported to influence blood composition^17,18^. Test statistics before (x-axis) and after adjustment (y-axis) for such nearby SNPs are all along the diagonal, indicating the reported SNPs do not confound the analysis. C) Correcting for latent factors leads to slightly more significant results. Depicted are the test-statistics in the original analysis, corrected for latent factors (x-axis), and the test-statistics without correction for these latent factors (y-axis). D) Multi-SNP GIs outperform single-SNP GIs in terms of predictive ability of index gene expression. The *F*-statistic calculated in the test set using the main, strongest associated SNP in the GIs is plotted against the *F*-statistic calculated using the full GI. Using the full GI results in 6,600 GIs predictive of the corresponding index gene (*F*-statistic > 10), whereas a single-SNP approach results in 4,910 predictive GIs.

**Figure S4.**
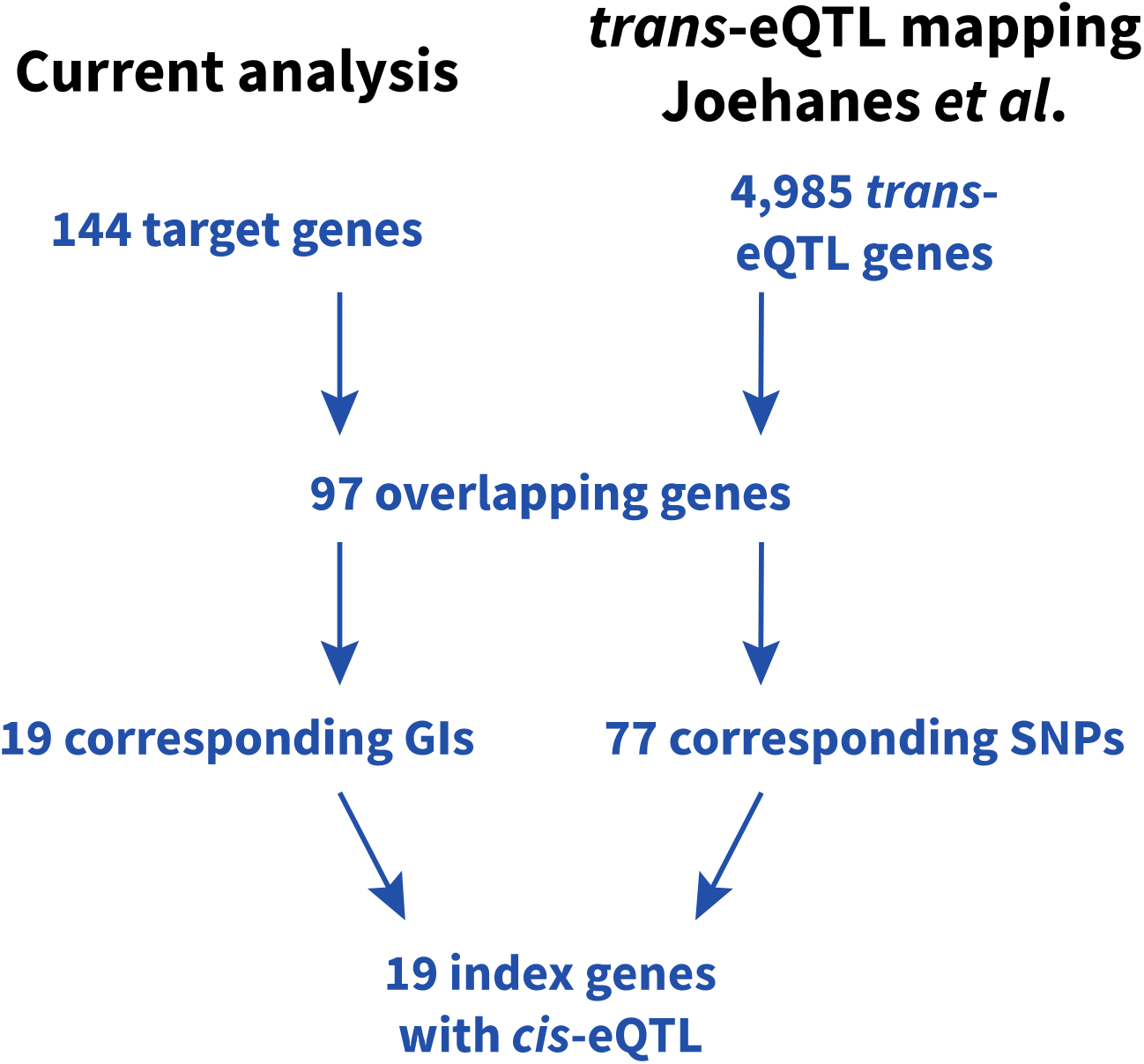
Diagram comparing the identified effects in the current analysis and those identified by an earlier *trans*-eQTL mapping effort^21^.

**Figure S5.**
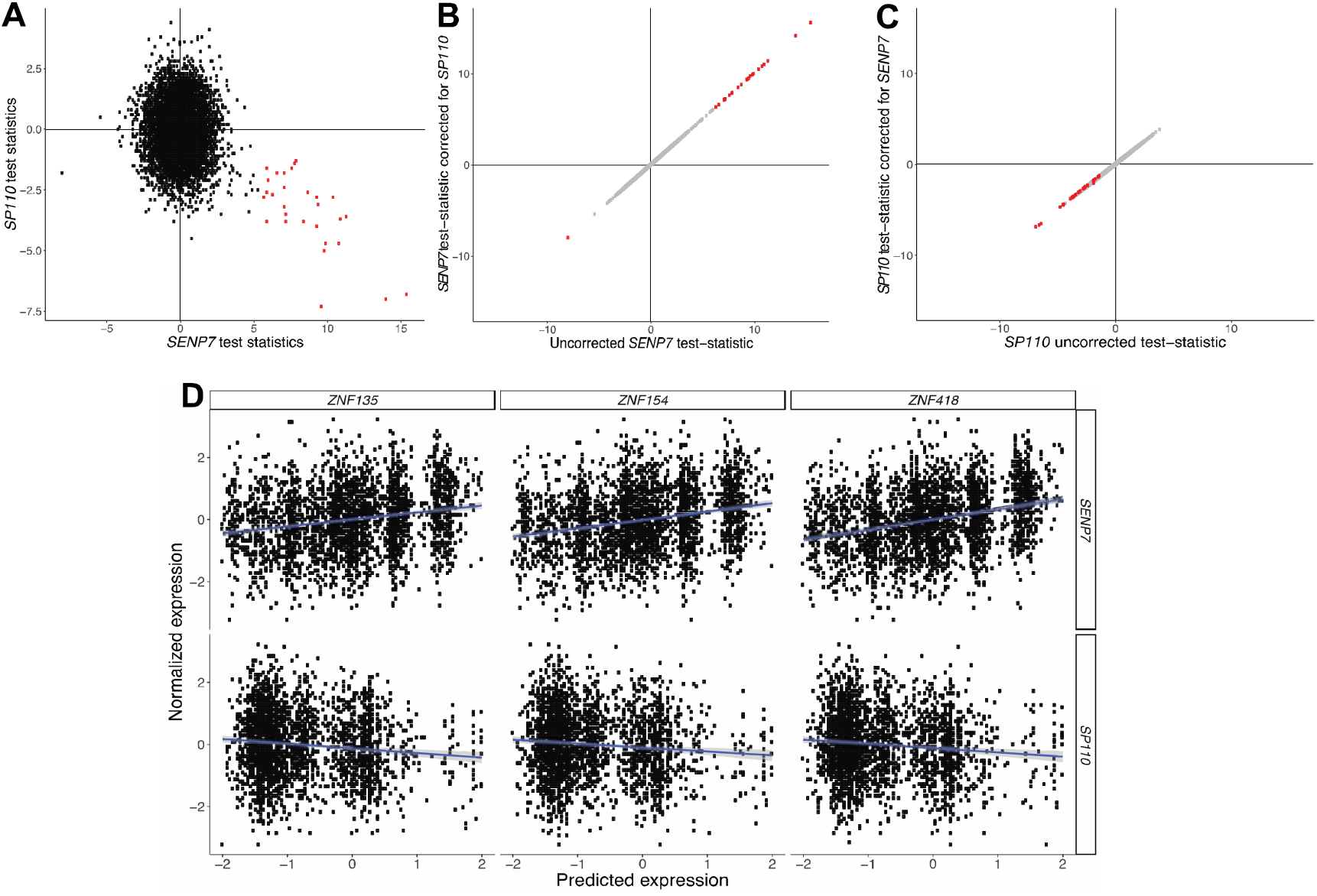
*SENP7*and *SP110* have shared, but opposite effects on the zinc finger protein cluster on chromosome 19. A) Test-statistics for *SENP7* and *SP110* show consistent opposite effects on the ZNF-cluster. B, C) Test-statistics of the directed effects of *SENP7* and *SP110* on target genes, correcting for each other’s genetic instruments (GIs). The unchanged test-statistics indicate their effects are independent. D) Illustrations of shared, but opposite effects.

